# Functional Screening of Human Small Intestinal Lactobacilli Reveals Strain-Specific Modulation of Epithelial Immune and Hormonal Responses

**DOI:** 10.64898/2026.08.04.742439

**Authors:** Erika J. Nachman, Lakshmi N. Somasundaram, A. Kyle B. Ardis, Sasirekha Ramani, Robert A. Britton

## Abstract

The human small intestine (SI) microbiome is a dynamic ecosystem that engages in complex interactions with the host, including those mediated by diet, hormones, and immune systems. However, the cultivation and characterization of small intestinal microbial therapeutics have yet to be developed. One reason for their limited progress may be that previous microbes were derived from other environments, including food, stool, and breast milk, rather than directly from their native environment, the human SI. Therefore, we investigated the human SI as a relevant source and vital target for the development of future microbial therapeutics for SI diseases. We isolated 10 unique *Lactobacillaceae* isolates from six different species from samples spanning the upper gastrointestinal tract of five organ donors. We performed whole-genome sequencing analysis and assessed the viability of isolates after numerous GI-like stresses. Next, we examined the ability of the isolates to stimulate enteric hormone secretion, modulate pro-inflammatory cytokines, and influence the replication of a live, attenuated oral rotavirus vaccine strain. We observed that SI *Lactobacillaceae* strains can survive GI-related stresses like other commercialized lactobacilli. Certain isolates were able to promote secretion of the hormones secretin and oxytocin from *ex-vivo* adult tissue and human infant intestinal organoids. In addition, we found strains that were able to modulate TNF secretion from the human monocytoid line THP-1. Finally, we found that *L. rhamnosus* strain 103 (Lr 103) significantly promoted antiviral IFN-λ secretion via TLR3 by secreted RNA in infant organoids. Lr 103, restricted rotavirus vaccine strain replication in infant organoids.

**Importance:** The impact of the microbiome on human health and disease has highlighted the potential of using microorganisms to prevent and treat human disease. The small intestine has been dramatically under sampled in regard to the intestinal microbiome. Here we isolate several novel organisms from the small intestine and focus on new isolates from lactic acid bacteria that are currently used as probiotics. We find that individual isolates have the ability to impact gut hormone secretion and modulate the immune system. Our work demonstrates the human SI as a relevant source of potential microbial therapeutics through the isolation of novel strains and their modulation of host physiology in preclinical models.

## Introduction

The human small intestine (SI) is a dynamic and critical site of physiology and host-microbe interactions where the SI epithelium hosts immune education, nutrient absorption, metabolic regulation, pathogen defense, and interactions with the microbiome^1,2,3,4^. Research over the last 20 years has revealed that the human gut microbiome influences immune development, maintains epithelial barrier function, synthesizes essential vitamins, and protects against enteric pathogens^5^. However, most of these research efforts have centered on large intestinal microbial communities, where sample accessibility and microbial density enabled deeper characterization, and the human SI microbiome and its impact on SI physiological remains comparatively understudied. This gap is particularly relevant given the millions of people with irritable bowel syndrome (IBS), Crohn’s disease, and other functional gastrointestinal disorders that amount to $358 million in healthcare direct costs^6^.

Microbes have been examined as potential treatments or preventatives to combat SI disease. Although microbial therapeutics such as VOWST and REBYOTA effectively prevent recurrent *Clostridioides difficile* in the large intestine, no Food and Drug Administration (FDA) approved microbial therapeutics exist for SI diseases. This may be in part because these microbes are frequently derived from stool, fermented foods, or non-intestinal sources such as breast milk, with limited relevance to the distinct environment of the SI. Previous candidate therapeutic microbes were also chosen based on taxonomy or historical use in food without a priority on functionality^7^. There is now increased recognition for the strain-level and niche-specific functions that could be implemented in the microbial therapeutic field ^8, 9,10^.

Given these gaps in microbial source and therapeutic target, the SI represents a favorable site for microbial therapeutic intervention. Enteric-coated ingested microbes reach the SI in higher concentrations than the large intestine when the microbial contents are released in the upper gastrointestinal tract (GIT)^11^. Once released in the SI, the microbial therapeutic can impact key physiological activities, including nutrient absorption, immune training, enteric hormone secretion, and response to live, oral vaccines^12,13^. However, the microbial therapeutic must adapt to the harsh environment of increased transit time, increased antimicrobial peptides, lower pH, exposure to bile, chyme, and pancreatic secretions. This intense SI environment shaped unique microbial communities with the increased presence of *Streptococcaceae*, *Lactobacillaceae*, and *Veillonellaceae* compared to the *Bacteroides*, *Lachnospiraceae*, *Ruminococcaceae*, and *Clostridiales* that dominate in the large intestine^14^. Given its distinct physiology, immune landscape, microbial composition, and expansive disease burden, we propose that the human SI represents an underexplored reservoir of microbes that can impact SI physiology and serve as candidate microbial therapeutics.

Here, we functionally screened novel *Lactobacillaceae* SI isolates for enteric hormone, immune, and enteric vaccine modulation. The majority of this study was conducted using an infant organoid model, as infants represent a vulnerable population with increased enteric infection burden, and the infant organoids are enriched in hormone-producing cells compared to adult organoids, allowing for the measurement of enteric hormones^15.16^. We observed that SI microbes were capable of modulating host responses, supporting further isolation and characterization of SI microbes and their potential impact on human health and disease.

## Methods

### Organ Inclusion and Exclusion Criteria

The inclusion and exclusion criteria for organ donations were reported in Nachman et al^17^. In brief, our inclusion criteria were donors older than 18 years of age, hospitalized in the Texas Medical Center, with no known bowel disease, surgery, or trauma (Inflammatory bowel disease, *Clostridioides difficile* infection), hospitalized for less than 28 days, and brain-dead (BD) for less than 7 days. We accepted both BD and donation after circulatory death (DCD) donors. DCD donors had a cut-off time of 90 minutes after the removal of life-support machinery for the inclusion of the organs in this study.

The exclusion criteria included patients with positive serological tests for hepatitis B, hepatitis C, or HIV, significant GIT bleeding during hospitalization, prior intestinal resection, a medical examiner case, transfer to the hospital from a long-term care facility, cases with GIT tissue recovery following organ recovery, or our lab being unable to process the specimen^17^. The organ processing has been described in Nachman et al., and extended demographic information on the organ donors can be found in **Table S1**^17^.

### Media compositions

Five different media types were used to plate the samples across all the donors **(Table S2).** Each media type used the base, supplement, and basal media but differed in the addition of mucin, bile, and short-chain fatty acids to create media to enhance GI microbial growth **(Table S2)**. Each sample was plated once on all media types described in **Table S2**.

### Culturing and purifying SI microbes

#### Mucosal sites

Mucosal scrapings from each segment of the intestinal tract were plated at 10^0^ and a 10^−1^ dilution in the microoxic chamber after processing^17^. Scrapings from the large intestine were plated anaerobically with and without the addition of reducing agents (0.4 g L-cysteine HCl, 1.6 g sodium thioglycolate, dithiothreitol, and 10 mL of sterile water), at 10^0^ and 10^−1^ dilutions in sterile phosphate-buffered saline (PBS).

#### Luminal Sites

The small intestine luminal samples were plated at 10^−1^ and 10^−2^ dilutions in the microoxic chamber, except for the stomach, which was plated at 10^0^ and at 10^−1^ dilutions after processing^17^. Luminal samples from the large intestine were plated in anaerobic conditions with and without reducing agents, where the ascending colon was plated at 10^−3^ and 10^−4^; the transverse colon was plated at 10^−4^ and 10^−5^, and the descending colon and appendix were plated at 10^−5^ and 10^−6^. Samples were plated on all the media types described in **Table S2**.

All samples were diluted in sterile PBS and incubated at 37°C for 24-72 hours. A subset of the resulting colonies was purified by streaking for isolation and stored for downstream sequencing and characterization. After colonies were picked to a new plate for purification, the remaining colonies on the plate were washed using sterile PBS and saved; these are referred to as “plate washes”. All samples saved were frozen stocks at −80°C in 1:1 in 30% PBS-glycerol.

### DNA isolation and V4 16S rRNA amplification of SI communities

During sample collection, luminal and mucosal samples were stored in 10% DMSO at −80°C for 16S rRNA sequencing. DNA from initial samples in addition to plate washes were extracted using the DNeasy Qiagen PowerSoil Kit Pro (#47014, Qiagen). The manufacturer’s protocol was followed with the addition of two 30-second bead beating steps with one 1-minute rest in between at the start of the extraction. The extracted DNA was eluted in 30-50 μL of elution buffer. The samples were submitted to the Alkek Center for Metagenomics and Microbiome Research CMMR facility, where library preparation and V4 sequencing were performed. Amplification of the V4 region was completed using 515F (5’-GTGCCAGCMGCCGCGGTAA-3’ and 806R (5’-GGACTACHVGGGTWTCTAAT-3’) with adapters for MiSeq and a single-index barcode in the reverse primer to later pool PCR products to 25,000 read pairs^161^. The raw data (BCL format) were then converted to FASTQ and demultiplexed from the single-index barcode in the Illumina bcl2fastq software. The demultiplexed reads were filtered for quality using bbduk.sh (BBMap, version 38.82)^19^, which removes adapters, PhiX reads, and low-quality Phred scores. Quality-controlled reads were then merged using bbmerge.sh with merge parameters: maxstrict=t, qtrim=t, trimq=15. Merged reads were filtered further with vsearch, utilizing parameters optimized for the appropriate 16S amplicon type^20^. The resulting reads were combined into a single FASTA for DEBLUR^21^. DEBLUR was set to a 252 bp limit, and the representative sequences were mapped against the most current SILVA database with a 97% cut-off for identity threshold^22^. Phylogeny information contained in the biom file is generated by aligning the centroid sequences with MAFFT^23^, and FastTree created a tree^24^. The biom file and the number of reads per sample are merged with a file generated for the overall read statistics to produce a final summary file with read statistics and taxonomy information.

### ATIMA analysis

The V4 16S rRNA sequencing was analyzed using webtool *Agile Toolkit for Incisive Microbial Analyses (*ATIMA). ATIMA used the R function vegan:adonis version 2.5.5 to estimate PERMANOVA p-values, and adjusted for multiple comparison p-values are adjusted for multiple comparisons with Benjamini and Hochberg’s formula to control for the false discovery rate^25,26^.

### Targeting the isolation of *Lactobacillaceae* from organ donors

100 µL of donor samples were spread plated on MRS (De Man–Rogosa–Sharpe, BD Difco) plates with 1.5% (w/v) agar, 300 µg/mL of vancomycin to selectively isolate *Lactobacillaceae* (J62790.03, Thermofisher) and 10 µg/mL of fluconazole (1271700, SigmaMillipore) or 2.5 µg/mL of amphotericin B (A2942, SigmaMillipore) to reduce GI fungi. These were incubated at 37°C in atmospheric oxygen conditions for 24-48 hours.

### Whole Genome Sequencing and Genome Assembly of SI *Lactobacillaceae*

Microbial isolates were sent to SeqCenter (Pittsburgh, PA) for 200 Mb Illumina whole-genome sequencing with 150 bp paired-end reads. Genomes were assembled using Unicycler (v0.5.0)^27^.

### Taxonomy Classification and Average Nucleotide Identity Analysis of SI *Lactobacillaceae*

Assembled genomes were compared to the Genome Taxonomy Database for taxonomic identification to the closest related species using the classify command^28^. To compute the average nucleotide identity, FAST_ANI maps each isolate in a pairwise manner and compares the ANI of mapped orthologs^29^. All bioinformatics work was conducted on a SLURM-based cluster managed by the Biostatistics and Informatics Shared Resource (BISR), supported by NCI P30-CA125123 and institutional funds from the Dan L. Duncan Comprehensive Cancer Center and Baylor College of Medicine.

### Bacterial supernatant production

SI *Lactobacillaceae* isolates were incubated for 18 hours in MRS broth. The following day, the culture was diluted 1:50 into LDM4^30^, RPMI-1640 (Sigma-Aldrich, R8758), or DMEM (Gibco, #11965092), as noted in the specific figure legends, until OD_600_ 0.5-0.6 was achieved. The cultures were centrifuged at 4000 rpm for 5 minutes (Thermo Fisher Scientific Heraeus Multifuge X3R Benchtop Centrifuge with a 75003180 rotor), and the supernatant was transferred to a new tube for neutralization using 0.1-10 M NaOH. The neutralized supernatant was sterilized by a 0.22 μM syringe filter, aliquoted, and stored at −20°C for up to 6 months.

### Bacterial isolates challenged with PAM3CSK4 on THP-1 cells

THP-1 cells were differentiated for 48 hours using 100 nM of phorbol 12-myristate 13-acetate (PMA) (P1585, Millipore Sigma). After differentiation, macrophage-like THP-1 cells were primed with 0.1 ng/μL of Pam3CSK4 (PAM) (Millipore Sigma, AABH9A954F84) in RPMI (Fisher Scientific, Corning, MT10041CM) overnight. The following day, the priming was removed, and 20% of bacterial supernatant and 80% RPMI were added with or without 1 ng/mL of PAM. The cells were treated for 4 hours, and ELISA (R&D DuoSet, DY210-05, DY201-05, and DY208-05) was completed to quantify cytokines.

### Organoid propagation and 2D monolayer formation

Infant organoids were purchased from the BCM organoid core, propagated in CMGF+ and incubated at 37°C, 5% CO_2_^31^. Jejunal (J) and ileal (L) organoid lines used were J1005, J1006, J1009, IL1002, IL1004, IL1010, and IL1013^15^. Demographic information on the organoid lines used is summarized in **Table S3**. Organoid propagation and monolayer methods were followed as according to published protocols^31,32^. In brief, infant organoids were plated as single cell suspension on Matrigel-coated 96 well plates and incubated in growth media (CMGF+) for 2 days^31,32^. After two days, the growth media was replaced with differentiation media for four days^31,32^. Monolayers were visually assessed by light microscopy after four days of differentiation for 95-100% confluency.

### SI *Lactobacillaceae* supernatant treatment on infant organoid to assess IFN-λ secretion

After complete removal of the differentiation media, 100 μL of the mid-log cell-free bacterial supernatants were applied to the monolayers for 24 hours with 100 μg/mL of polyI:C (polyI:C hmw; Invivogen) as the positive control at 37°C, 5% CO_2_.

PolyI:C priming experiments were performed by treating the cells with a lower dose of polyI:C (50 μg/mL) as the primer, diluted in DMEM for 4.5 hours. The polyI:C treatment was completely removed, and then the cells received 100 μL of bacterial supernatants, DMEM media control, or polyI:C (100μg/mL) in DMEM as the positive control for 18 hours at 37°C, 5% CO_2_. IFN-λ was measured by ELISA (DY1598B, Human IL-29/IL-28B DuoSet ELISA).

### Inhibition of TLR signaling on infant organoids

To assess the effects of TLR signaling, differentiated infant organoid 2D monolayers were first primed with 50 μg/mL of polyI:C in DMEM for 4.5 hours. The polyI:C prime was discarded, and the organoids were treated with 100 μL of Lr 103 supernatant, 100 μg/mL of polyI:C in DMEM, La 180 supernatant, and DMEM with or without 500 mM of FC-99 hydrochloride (SML2371, Sigma-Aldrich) and 200 μM of TL2-C29 (inh-c29, Invivogen) for 18 hours at 37°C, 5% CO_2_. J1005 was used across 3 independent experiments, and IFN-λ was measured by ELISA.

### RNase Treatment on Infant Organoids

The dsRNA in the supernatant was depleted by the addition of 15 μL of RNase III (AM2290, Invitrogen) and 5 μL of reaction buffer to 700 μL of Lr 103, polyI:C (100 μg/mL), and DMEM. These reactions were incubated at 37°C for 1 hour. For ssRNAse treatment, 0.4 μL of RNase T1 (EN0541, ThermoScientific) was used to treat 700 μL of supernatant or controls and incubated for 30 minutes at 37°C. The RNAses were inactivated through incubation at 70°C for 10 minutes. In parallel, J1005 differentiated infant monolayers were primed with 50 μg/mL of polyI:C in DMEM for 1.5 hours. The ssRNAse, dsRNAse treated supernatants and controls were then applied to the organoids in addition to no treatment controls and incubated at 37°C, 5% CO_2_ for 18 hours, and IFN-λ was measured by ELISA.

### Rotavirus Vaccine Infections

Ileal and jejunal infant organoids were plated on transwells (Corning #3415) coated with collagen IV (Sigma-Aldrich, C5533) at 2.5 −10^5^ cells/mL. Following 4 days of differentiation using an in-house differentiation media^15^, the cells were apically treated with 200 μL of mid-log cell-free bacterial supernatant, DMEM as a negative control, or 100 μg/mL of polyI:C as a positive control for 18 hours at 37°C, 5% CO_2_. The next day, the cells were then infected basolaterally with lab-adapted Rotarix^TM^ vaccine (also called RV1) that was activated with 10 μg/mL of Worthington trypsin at an MOI of 0.5 for 2 hours at 37°C, 5% CO_2_. Apical treatments were not removed during the 2 hours of virus inoculation. At the end of 2 hours, the apical media was collected and the trans-wells were washed twice with CMGF media to remove any unbound virus. The trans-wells were then placed in 500 μL of differentiation media while the apical side received 200 μL of mid-log cell-free bacterial supernatant, DMEM or polyI:C as added in the previous step. The organoids are incubated at 37°C, 5% CO_2_ for a total of 24 hours. Virus titers in the apical samples at 2 hours and 24 hours post infection were tittered using a Fluorescence Focus Assay (FFA) on MA104 cells as described previously^33^. Briefly, 3-fold dilutions of trypsin activated apical samples were added to confluent MA104 cells and were incubated at 37°C, 5% CO_2_ for 1 hour. The cells were then washed, 100 μL of DMEM media was added to each well and the cells were incubated at 37°C, 5% CO_2_ for an additional 15 hours. Following incubation, the cells were fixed with ice-cold methanol and stained with an in-house rabbit anti-rotavirus primary antibody and a donkey anti-Rabbit Secondary Antibody (IgG (H+L) Highly Cross-Adsorbed, Alexa Fluor™ 488, Invitrogen). A Biotek Cytation 7 cell imaging reader was used to count the number of infected cells per well. IFN-λ levels were tested in the basolateral samples by ELISA.

**Figure 1.**
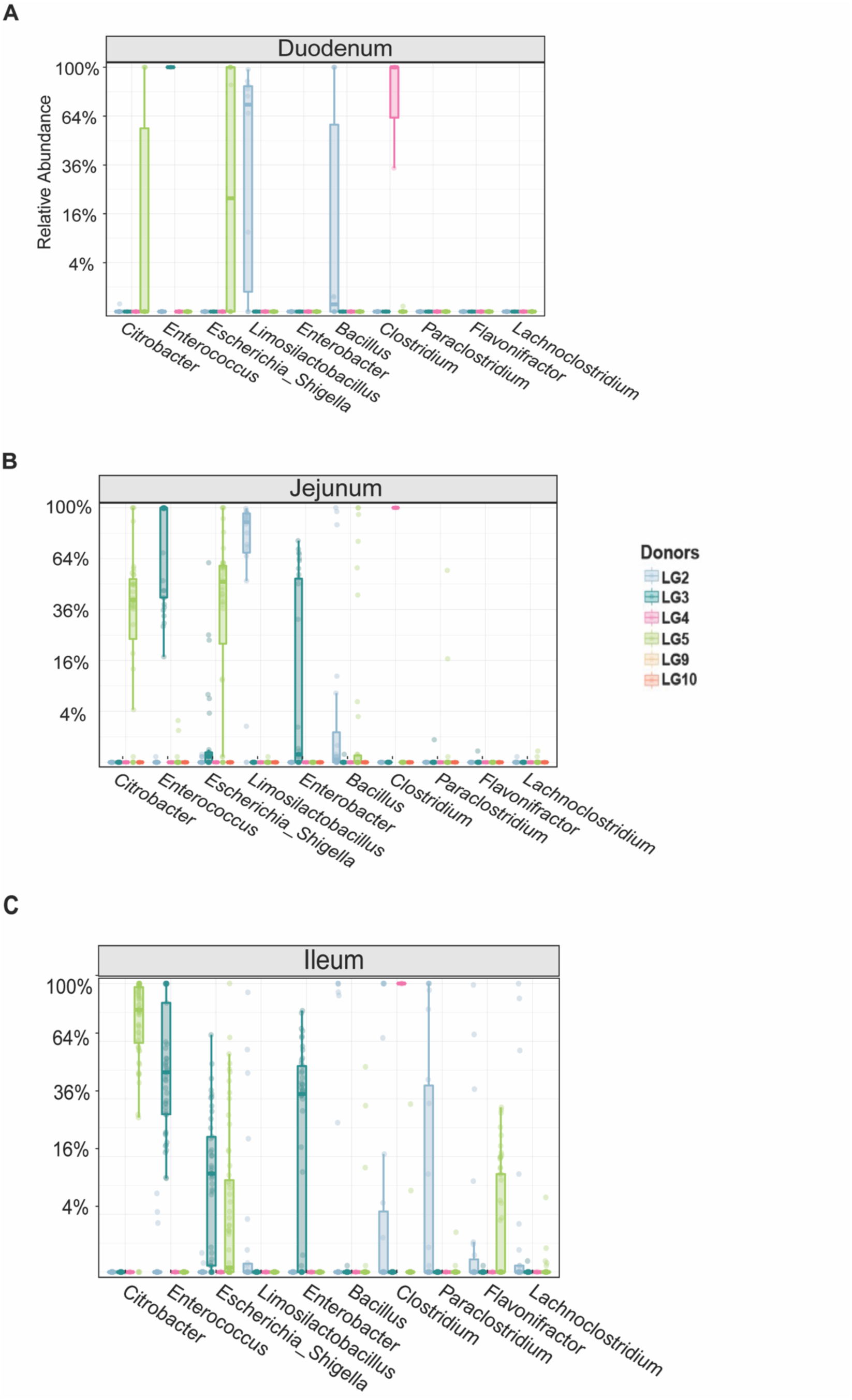
Cultivation of SI microbial community from organ donors. The top 10 taxa are plotted for each cultivated sample from the duodenum **(A)**, jejunum **(B)**, and ileum **(C)** for donors 2-5, 8, and 10. Cultivated samples summarize the cultivation efforts from plate washes of bacterial communities grown under both microoxic and anaerobic conditions.

### Toxicity of Bacterial Supernatant on Infant Organoids

To ensure that the organoids were viable after experimentation with bacterial supernatants, PrestoBlue (A13261, ThermoFisher) assays were performed according to the manufacturer’s protocol as referenced in the specific figure legends. In brief, 1X sterile PBS-PrestoBlue solution was treated on the infant organoids for 20 minutes at 37°C, 5% CO_2_. A minimum of two wells were treated with 70% ethanol for 5 minutes prior to the assay as a control for complete lysis. Alternatively, CellTox Green Cytotoxicity Assay was performed on uninfected organoids apically treated with DMEM, polyI:C, or the bacterial supernatants for 24 hours according to the manufacturer’s protocol with a well dedicated as a complete lysis control **(Fig. 6)** (G8731, Promega).

### Statistical Analysis

All statistical analyses were performed in R (version 4.5.2, 2025-10-31). Data in Figures 2–6 were analyzed using linear models or linear mixed-effects models, as appropriate. In **Fig. 2**, the hormone data were analyzed using treatment as a fixed effect with donor or experiment as a random intercept. Cytokine data in **Fig. 3** were analyzed using a linear model with treatment as a fixed effect, with each measurement treated as an independent observation. **Fig. 4 and Fig. 5** were analyzed with a linear mixed-effects model with the experiment included as a random effect and treatment as the fixed effect. Viral replication data in **Fig. 6** were analyzed with treatment condition as fixed effects, and experiment as a random intercept. Viral loads were log_10_-transformed prior to analysis. For all data, emmeans was used to calculate the marginal means and complete pairwise comparisons. Multiple comparison significance values were adjusted by Benjamini–Hochberg.

**Figure 2.**
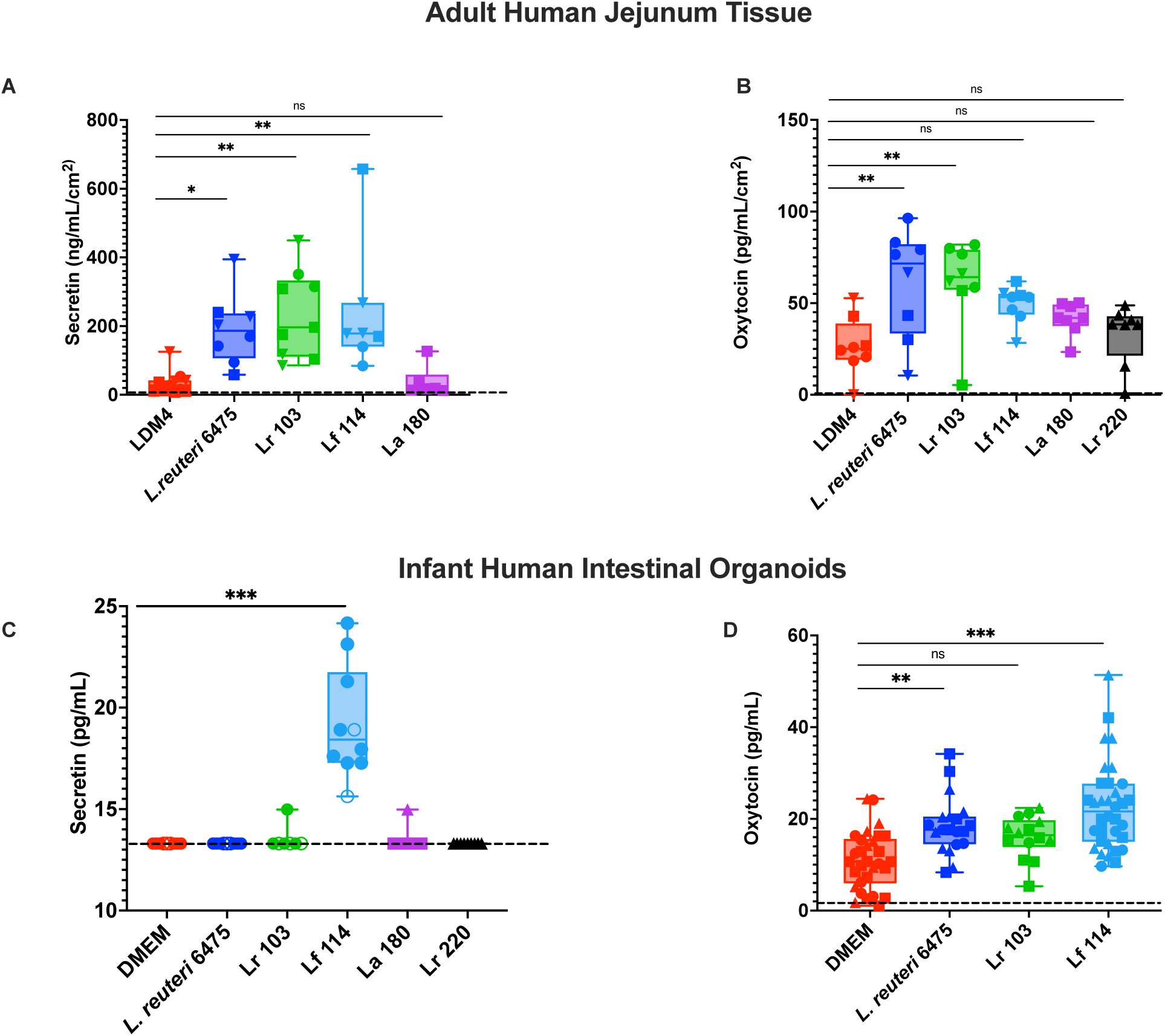
Lr 103 and Lf 114 can stimulate secretin and oxytocin from SI intestinal models. Tissue sections were bathed in 3-5 mL of bacterial supernatant grown in Lactobacillus defined media 4 (LDM4) for 3 hours, and an ELISA was completed to measure secretin **(A)** or oxytocin **(B)** (n=3). The SI-isolated bacteria examined were *L. rhamnosus* 103 (Lr 103), *L. fermentum* 114 (Lf 114), *L. animalis* 180 (La 180), and *L. rhamnosus* 220. **(A-B)** Each shape represents an independent organ donor tissue sample. **(C)** Bacterial strains were grown in LDM4, and the resulting cell-free bacterial supernatant was treated on jejunal infant organoid differentiated monolayers J1005 (closed circle), J1006 (triangle), J1009 (open circle) to measure secreted secretin by Luminex (3 organoid lines, 2 independent experiments, 2 technical replicates). **(D)** Cell-free bacterial supernatant was produced in DMEM and treated on jejunal and ileal infant organoid differentiated monolayers J1005 (closed circle, n=4 experiments, 2 technical replicates) J1006 (triangle, closed circle, n=4 experiments, 2 technical replicates), IL1002 (square, n=7 experiments, 2 technical replicates). Statistics were completed using a mixed linear model where the treatment was a fixed effect with donor or experiment as a random intercept. Multiple comparison significance values were adjusted by Benjamini–Hochberg. The dotted lines represent the limit of detection. Significances are designated by * indicating p ≤0.05, ** indicating p ≤0.01, and *** indicating p ≤0.001.

**Figure 3.**
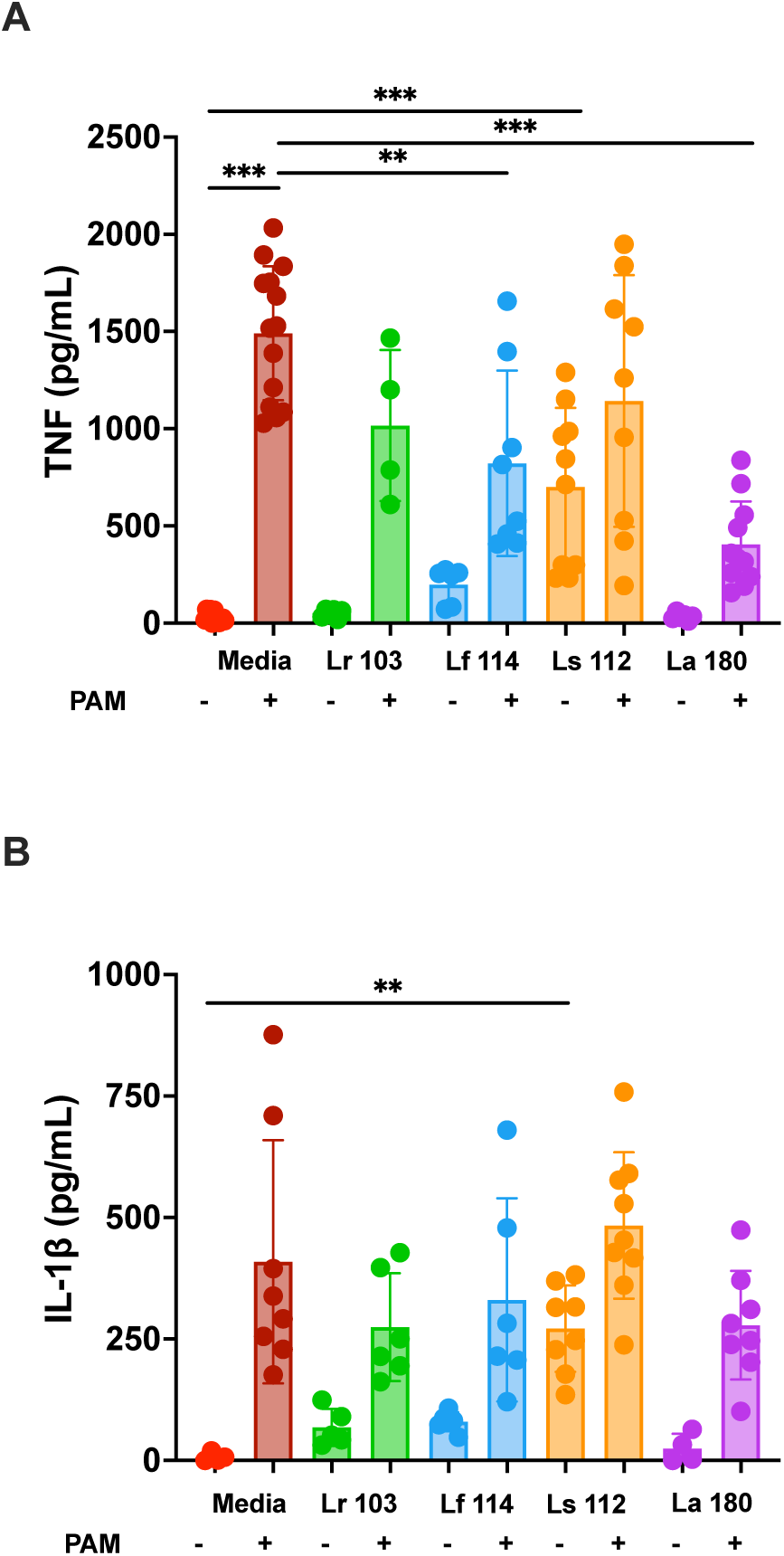
La 180 can decrease TNF-alpha in macrophage-like cells after TLR2 agonist priming. Bacterial supernatants were added to THP-1 cells or primed with TLR-2 agonist, and cytokines TNF**(A)** and IL-1β **(B)** were measured using ELISA (n=4-6). Statistics were calculated by a linear model with treatment as a fixed effect and each measurement treated as an independent observation. Multiple comparison significance values were adjusted by Benjamini–Hochberg. Significance is designated by * indicating p ≤0.05, ** indicating p ≤0.01, and *** indicating p ≤0.001.

**Figure 4.**
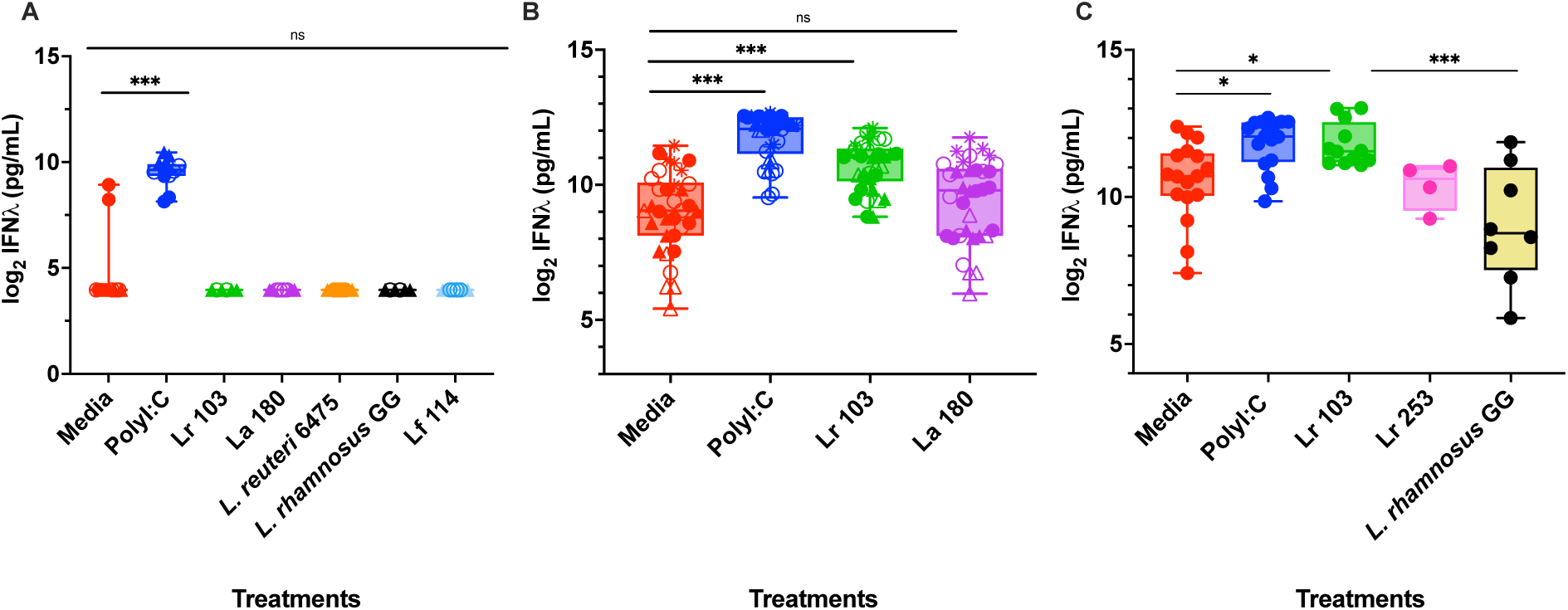
Lr 103 promotes release of IFN-λ from infant intestinal organoids. (**A**) Addition of bacterial supernatant alone on infant monolayers does not elicit secretion of IFN-λ (n=2 independent experiments. J1005 is depicted as a closed circle (n=3, 2 technical replicates). J1006 is displayed as a closed triangle (n=4, 2 technical replicates), and IL1002 is open circle (n=2, 2 technical replicates) **(B-C)** Lr 103 significantly promotes IFN-λ after polyI:C priming in all conditions on jejunal (J1005 open circle, n=9, J1006 open triangle, n=7) and ileal infant organoids (IL1002 open circle, n=8, IL1004 open triangle, n=5, IL1013 asterisks, n=5). Each point represents an average of the technical replicates per experiment. Data were analyzed using a linear mixed-effects model, with experiment included as a random effect and treatment as a fixed effect. Multiple-comparison significance values were adjusted using the Benjamini–Hochberg procedure. Significance is designated by * indicating p ≤0.05, ** indicating p ≤0.01, and *** indicating p ≤0.001.

**Figure 5.**
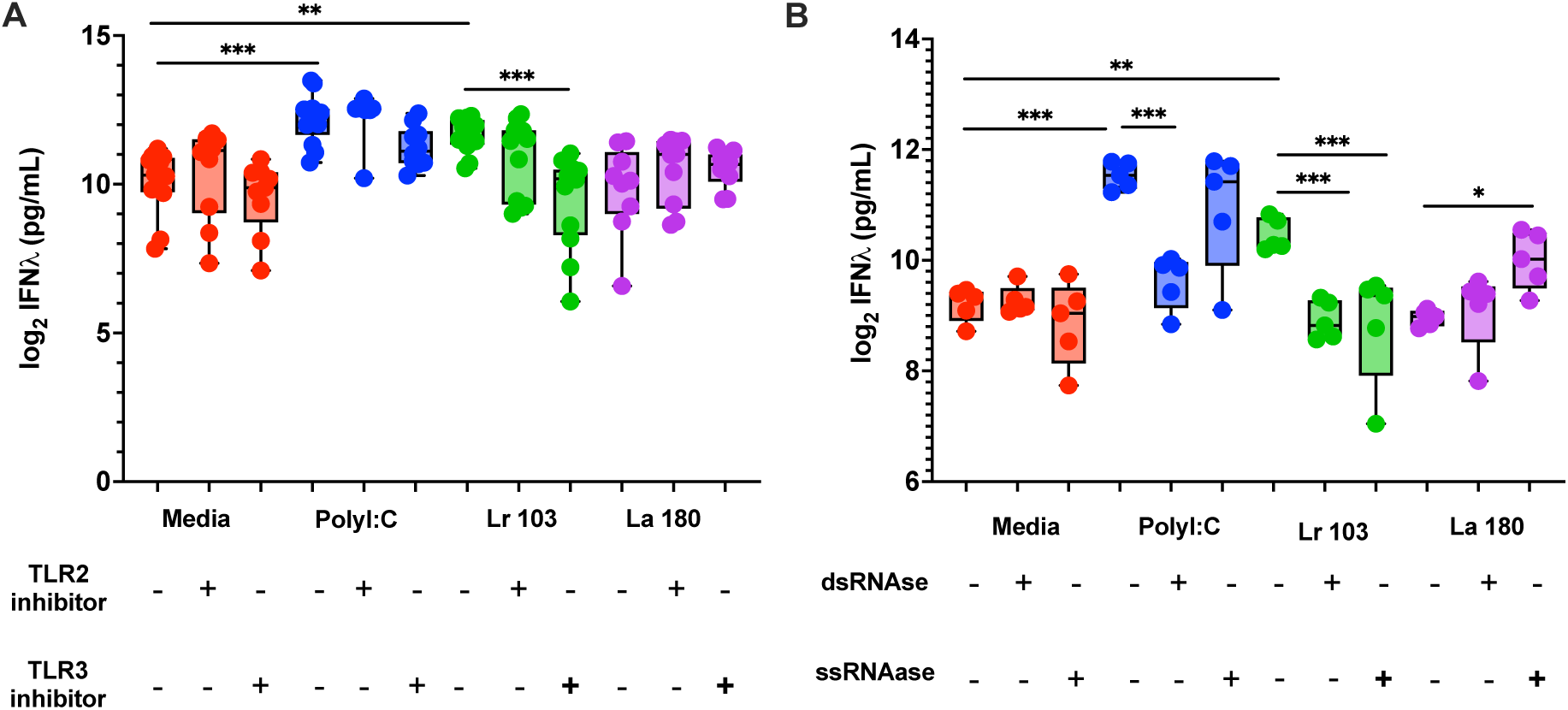
IFN-λ secretion is dependent on TLR signaling from a secreted RNA. **(A)** J1005 infant organoids were treated with TLR3 inhibitor, FC-99, and TLR2 inhibitor, C-29, at 500 mM and 200 μM, respectively (n=7, each point represents an average of 3 technical replicates per experiment). **(B)** dsRNAse and ssRNAse were treated on bacterial supernatants and transferred after inactivation to J1005 organoid monolayers for 3 hours (n=5, each point represents an average of 3 technical replicates per experiment). Data were analyzed by a linear mixed-effects model with the experiment included as a random effect and treatment as the fixed effect. Multiple-comparison significance values were adjusted using the Benjamini–Hochberg procedure. Significance is designated by * indicating p ≤0.05, ** indicating p ≤0.01, and *** indicating p ≤0.001.

### Data Availability

The 16S rRNA community data from **Fig 1**. And **Fig. S1**. can be found at: https://atima.research.bcm.edu/?p=6688&k=yY6hptYpjoXW8.Wlh3ed_A--. Whole genome sequencing files of *Lactobacillaceae* isolates the can be found at PRJNA1422122. R scripts can be found at https://github.com/erikanachman/statistics_2026/blob/main/.github/workflows/main.yml

## Results

### Isolation of microbes from the human small intestine

To start identifying and collecting small intestinal (SI) microbial isolates, we first performed sequencing on our samples that were saved directly from the donors, known as our culture-independent samples. Our cultivation-independent samples revealed that the SI harbored *Bacteroides*, *Streptococcus*, *Prevotella*, *Paraclostridium,* and *Escherichia/Shigella* in high relative abundance across the SI in multiple donors **(Fig. S1)**. Notably, several of our SI samples failed culture-independent sequencing (7/12) and did not produce any CFU after plating (2/12), indicating low biomass in the SI.

Culture-dependent efforts, the community of microbes washed off from the agar plate after culturing, resulted in the cultivation of several genera including *Escherichia* sp*., Lacticaseibacillus* sp.*, Enterococcus, Enterobacter, and Citro*bacter from duodenum, jejunal, and ileal samples grown under microoxic and anaerobic conditions and several types of culture media **(Fig.1)**. Interestingly, we observed genera that were only present from cultivation, such as *Flavonifractor*, *Enterobacter*, and *Citrobacter,* suggesting either that those genera were contaminants or that a select few low-abundance members outcompeted others and grew well under the conditions. Nevertheless, we cultivated several SI-associated microbes and began collecting isolates that can be further examined for potential biotherapeutic targets **(Fig.1)**. The *Lactobacillaceae* family was a promising target for further isolation due to their extensive history of beneficial properties and current commercial use as probiotics^34^.

### Isolation of *Lactobacillaceae* from the human SI

Culture-independent sequencing and cultivation data were used to target *Lactobacillaceae* from SI samples **(Fig. 1, Fig. S1)**. With this approach, we isolated 21 *Lactobacillaceae* from 6 individuals **(Table 1)**. However, we isolated the same species multiple times from different samples from the same donor for *Lacticaseibacillus rhamnosus* (9)*, Lacticaseibacillus paracasei* (5)*, and Limosilactobacillus fermentum* (4). Average nucleotide identity analysis revealed that the isolates from the same donor were 99.7-99.9% similar to each other for the 3 species, with one exception of *L. paracasei* 123 (98.8%), indicating high similarity and isolation of the same strain from multiple samples **(Fig. S2A-C)***, Ligilactobacillus animalis, Ligilactobacillus salivariusand Companilactobacillus nurkuri* were independently isolated once, totaling 10 unique isolates from 6 different species **(Table 1).**

**Table 1.** Lactobacilli isolated from the human upper gastrointestinal tract of organ donors. . Species-level identification of isolates recovered from independent donors. Each isolate name is an arbitrary designation assigned for strain stocking and is used to identify the strain in subsequent figures.

| **Species** | **Isolate name** | **Donor** | **Body Site** |
| --- | --- | --- | --- |
| *Lacticaseibacillus*<br>*rhamnosus* | 103, 104, 105,<br>106, 107 | 1 | Lower Ileum, Duodenum |
| *Lacticaseibacillus*<br>*rhamnosus* | 111 | 2 | Stomach |
| *Lacticaseibacillus*<br>*rhamnosus* | 220 | 5 | Middle Ileum |
| *Lacticaseibacillus*<br>*rhamnosus* | 236 | 10 | Duodenum |
| *Lacticaseibacillus*<br>*rhamnosus* | 253 | 12 | Duodenum |
| *Lacticaseibacillus*<br>*paracasei* | 108, 109, 110,<br>117 123 | 2 | Duodenum, Upper Jejunum,<br>Stomach |
| *Ligilactobacillus*<br>*salivarius* | 112 | 2 | Stomach |
| *Limosilactobacillus*<br>*fermentum* | 114, 115, 116,<br>119 | 2 | Duodenum, Middle Jejunum,<br>Upper Jejunum |
| *Ligilactobacillus*<br>*animalis* | 180 | 3 | Small Intestine |
| *Companilactobacillus*<br>*nurkuri* | 232 | 8 | Duodenum |

As we isolated multiple, independent strains of *L. rhamnosus* (Lr) from the donors, we examined whether the isolates were consumed from a commercial probiotic. We compared the average nucleotide identity (ANI) of strains Lr 103, Lr 111, Lr 220, Lr 236, and Lr 253 against commercial products or potential probiotic strains *L. rhamnosus* GG, *L. rhamnosus* BFE5264, *L. rhamnosus* 4B15, *L. rhamnosus* LDTM7511, and *L. rhamnosus* LKF7. The commercial strains were more similar to each other than the SI human isolated strains **(Fig. S2D)**. While this is not an exhaustive list of all commercial probiotics, these data suggested that these strains were not ingested as probiotics **(Fig. S2D)**. We additionally performed pairwise comparisons of the isolates’ average nucleotide identity (ANI) against their respective species type-strains **(Fig. S2E).** All strains were more than 97% identical to the type-strain, as defined for bacterial species similarity **(Fig. S2E).** Strains La 180 and Nc 232 were extremely similar to their respective type-strain (99.2%), which demonstrates high similarity in their core genomes **(Fig. S2E)**.

### SI *Lactobacillaceae* Grow Robustly and Resist Stress Conditions

The initial criteria for evaluating potential microbial therapeutics include robust growth under laboratory conditions, tolerance to GI-like stress, and demonstrated safety. These are a few of the criteria evaluated for potential probiotics. The *Lactobacillaceae* SI isolates grew well in general laboratory settings, and 7/8 isolates exceeded survival of exposure to pH 2 for 2 hours compared to probiotic *L. reuteri* 6475, ranging from 31-83% survival **(Fig. S3A-B).** The isolates were grown in increasing amounts of porcine bile, and we observed the isolates were able to maintain growth, albeit at a slower rate compared to control, demonstrating tolerance to bile exposure **(Fig. S3C-I).** Lastly, most of the isolates were susceptible to several classes of antibiotics **(Fig. S3J),** and none exhibited cytolytic alpha hemolysis activity **(Fig. S3K)**, highlighting their favorable safety profiles.

### Examining secretion of oxytocin and secretion from infant organoids and adult jejunal tissue

To examine relevant host-microbe interactions, we first examined the ability of the *Lactobacillaceae* isolates to impact enteric hormone secretion from infant organoids. Enteric hormones play important regulatory roles in the GI tract, and there has been increasing evidence of GI microbes altering enteric hormone secretion^35, 36, 37,38, 39^. For example, Danhof et al. demonstrated that *Limosilactobacillus reuteri* 6475 promoted oxytocin release from an enterocyte in a secretin-dependent manner in human gastrointestinal tract cells^30,40^. Therefore, we sought to determine whether SI-isolated microbes can alter secretin and oxytocin in similar models. We observed strain Lr 103 (*L. rhamnosus* strain 103) significantly promoted secretin and oxytocin up to 2.5-fold higher than media treatment on adult jejunal tissue, while other SI isolated microbes did not **(Fig. 2A, 2B).** While Lf 114 promoted secretin release from adult jejunal tissue, we observed a 2-fold increase in oxytocin release, although not statistically significant **(Fig. 2A)**. However, after treatment on infant organoids, only Lf 114 promoted secretin and oxytocin secretion **(Fig. 2C, 2D)**. These data suggest a broad induction of secretin and oxytocin by Lf 114, with a more context-dependent stimulation from Lr 103 and highlight species-specific responses within each model **(Fig. 2).**

### Assessing anti-inflammatory properties on THP-1 cells

The SI is a major site of immune training and is consistently exposed to potential pathogens^1,41^. Microbes have been examined as a tool to modulate inflammatory responses^34^. For instance, histamine produced by *L. reuteri* 6475 was shown to be anti-inflammatory by reducing TNF secretion from monocytoid cell line, THP-1 ^42^. Additionally, *L. rhamnosus* has been reported to promote reactive oxygen species that reduce pro-inflammatory responses in an immature intestinal mouse epithelium model^43^. Thus, we next examined the immunomodulatory activity of the SI *Lactobacillaceae* isolates. First, baseline TNF and IL1-β expression levels were established by using cell-free bacterial supernatants to treat the macrophage-like differentiated THP-1 cells **(Fig. 3)**. Our results indicated a pro-inflammatory response from Ls 112 supernatant, highlighted by significant secretion of TNF (713 pg/mL) and of IL1-β (271 pg/mL) on average above media control **(Fig. 3A, 3B)**. All other lactobacilli tested did not significantly induce either cytokine **(Fig. 3, Fig. S4A)**.

We next asked if the isolates could mitigate the effects of pro-inflammatory molecules such as the TLR2 agonist, PAM3CSK4 (PAM). We observed that after challenge with PAM on THP-1 cells, La 180 (*Ligilactobacillus animalis* strain 180) and Lf 114 significantly decreased TNF secretion by 3-fold and 2-fold, respectively, indicating a selective reduction of TNF by these strains **(Fig. 3A-3B).** SI microbes did not reduce IL-1β after PAM challenge. Furthermore, IL-8 secretion was found not to be promoted by SI nor mitigated by SI microbes **(Fig. S4A).**

### The secretion of antiviral IFN-λ from SI *Lactobacillaceae*

We next focused our investigation on whether these isolates influence epithelial-specific inflammatory pathways and provide protection against enteric pathogens. Enteric viral infections have a high mortality and case burden in infant populations, which could present a promising target for microbial therapeutic intervention^16^. Furthermore, previous work has demonstrated that members of the *Lactobacillaceae* family can elicit antiviral responses or decrease rotavirus replication in the gut^44^. We hypothesized that our strains may enhance epithelial antiviral defenses in infant models. A key mechanism of antiviral defense involves the production of interferons, including IFN-α, IFN-β, and IFN-λ, which promote expression of interferon-stimulated genes (ISG)^45^. Multiple studies have demonstrated that IFN-λ is the primary interferon secreted in the GI epithelium^46^ with modulation of homeostatic IFN-λ by the gut microbiome^47^. Therefore, we focused on an IFN-λ response after SI microbe treatment on infant organoids.

First, we aimed to understand the conditions under which IFN-λ is secreted in infant organoids. Upon stimulation with SI *Lactobacillaceae* isolate bacterial supernatants or polyI:C, a dsRNA synthetic analog known to stimulate IFN-λ through TLR3^48^, we observed a significant increase in IFN-λ secretion in only with polyI:C addition. This suggested that while IFN-λ is measurable in infant organoids, the bacterial supernatants alone were insufficient to promote IFN-λ secretion **(Fig. 4A)**.

We hypothesized that bacterial supernatants may enhance IFN-λ after a priming event with polyI:C. An initial screen revealed that Lr 103 induced IFN-λ secretion 3-fold above media alone across both organoid lines while La 180 did not **(Fig. S4B)**. The polyI:C priming experiment was repeated across 2 jejunal and 4 ileal infant organoid lines over multiple bacterial supernatant batches **(Fig. 4B)**. In agreement with the initial screen, Lr 103 significantly promoted IFN-λ over the media in ileal and jejunal organoids by 4-fold, capturing a consistent signal within the biological variation between the organoid lines **(Fig. 4B)**. We also observed that the promotion of IFN-λ was specific to Lr 103 while the other SI *L. rhamnosus* strain, Lr 256, and the probiotic *L. rhamnosus* GG did not significantly promote IFN-λ (**Fig. 4C**).

### Lr 103 promotes IFN-λ secretion through TLR3

Next, we investigated the mechanism by which Lr 103 increased IFN-λ secretion from the infant intestinal organoids. We hypothesized that this effect was through toll-like receptors on the gut epithelium, which detect a wide range of microbial ligands that induce specific immune responses^49^. We focused on TLR2, which recognizes bacterial acylglycerols, proteins, and polysaccharides^49^, and TLR3, which is known to respond to viral dsRNA^49^. To test their involvement, we added the TLR2 inhibitor TL2-C29 or the TLR3 inhibitor FC-99 to the supernatant treatments after polyI:C priming, then proceeded to measure IFN-λ secretion from infant organoids^50^. We observed that the addition of the TLR3 inhibitor significantly decreased IFN-λ secretion stimulated by Lr 103 supernatant in both organoid lines by 4-fold and 2.8-fold in the polyI:C conditions, whereas treatment with TL2-C29 had no effect **(Fig. 5A)**. FC-99 has previously been shown to downregulate TLR3 expression through an IRF3/IFN-a/JAK/STAT1 pathway and reduce systemic inflammation in a mouse model of sepsis^50^. These findings suggest that strain Lr 103 activates IFN-λ production through activation of TLR3 or engages pathways that converge on similar downstream targets of TLR3 **(Fig. 5A)**.

### IFN-λ secretion from Lr 103 is dependent on a secreted RNA

To investigate the role of TLR3 in our model, we examined whether strain Lr103 produces an RNA that could directly interact with TLR3. Previous studies have reported involvement of RNA in the promotion of interferons after bacterial supernatant^51,52^. Bacterial supernatants and controls were treated with an ssRNAse or dsRNAase, which were then incubated on infant organoids. We found that Lr 103 ssRNAse and dsRNAse-treated supernatants and polyI:C dsRNAse-treated supernatants significantly decreased IFN-λ secretion up to 4-fold **(Fig. 5B)**. This data suggests Lr 103 produced a secreted RNA that mediates the release of IFN-λ in the infant intestinal organoids **(Fig. 5B).** To ensure these effects are due to the bacterial supernatants, we performed cell viability and toxicity assays and observed the organoids remain metabolically active after treatment with the bacterial supernatants, inhibitors, and RNases **(Fig. S5A,5B).**

### Lr 103 reduces rotavirus replication in infant organoids

As IFN-λ possesses antiviral activity, we were then interested in whether Lr 103 could impact viral enteric replication *in vitro*^53^. Rotavirus, an enteric virus, is one of the leading causes of severe dehydrating gastroenteritis in infants in low-and middle-income countries despite available vaccines^54^. Previous work in adult intestinal organoids found that exogenous IFN-λ can reduce rotavirus titer by approximately tenfold ^55^. As children under 5 years are the most susceptible to rotavirus infection and the oral vaccine is administered in the first 6 months of life, infant organoids represent the most physiologically relevant model in which to examine the effect of bacterial supernatant on the live attenuated vaccine RV1 ^55^. We found Lr 103 restricted live-attenuated oral rotavirus RV1 replication by 0.6 log_10_ fold compared to media control **(Fig. 6A, Fig. S5C)**. La 180 also reduced rotavirus replication by 0.55 log_10_ fold on average, suggesting a general antiviral effect of *Lactobacillaceae* with a more consistent reduction by Lr 103.

**Figure 6.**
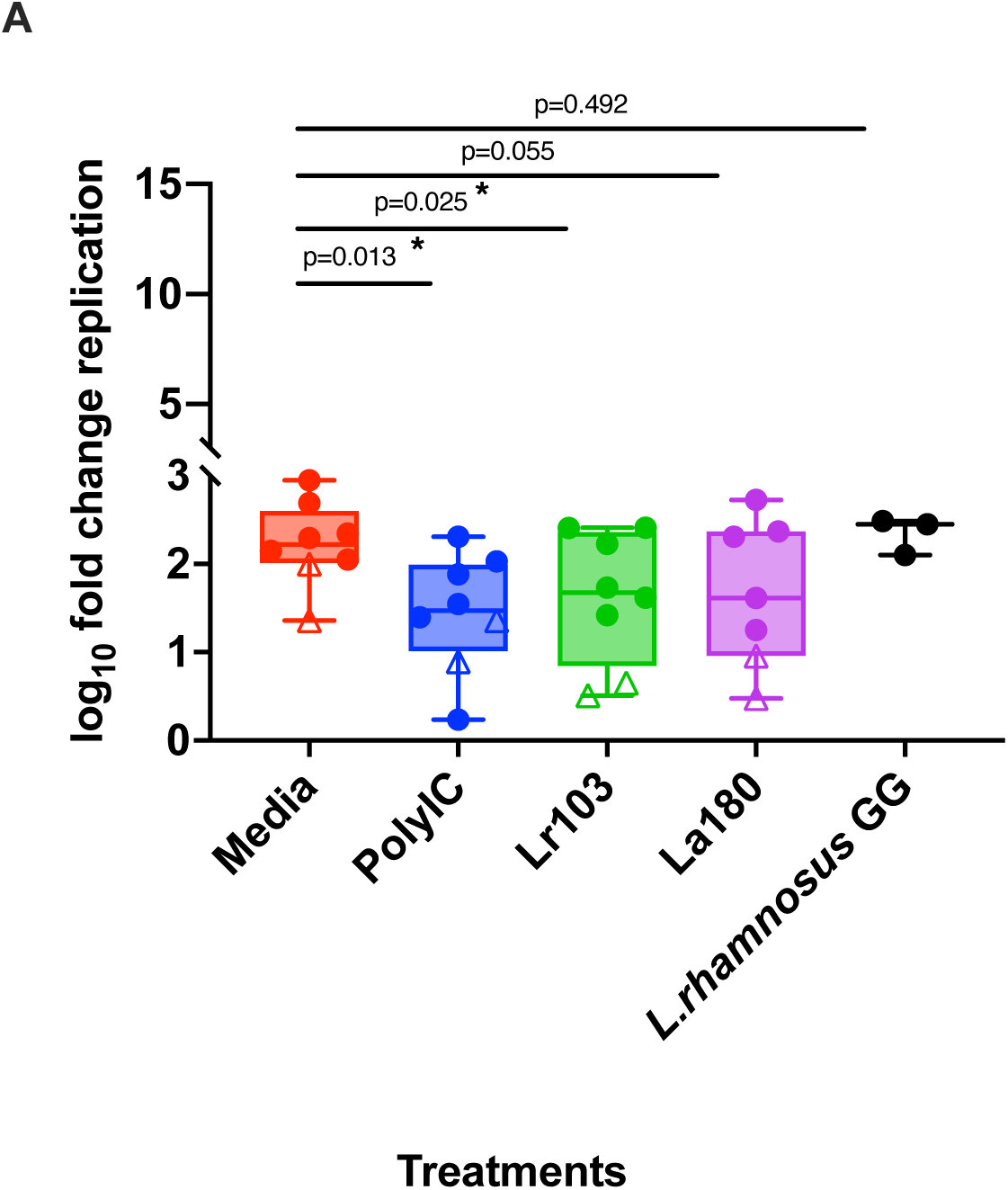
Lr 103 decreased rotavirus vaccine replication in infant organoids. **(A)** Lr 103, La180, polyI:C, *L. rhamnosus* GG, and media (DMEM) were treated on infected J1005 and IL1004 infant organoid trans-wells and infected with an MOI of 0.5 of RV1. RV1 replication is shown as log_10_ fold change, calculated by dividing viral RNA titers at 22 hours post infection (hpi) by the titers at 2 hpi. Each shape is an independent experiment. Significance was determined by a mixed linear model where treatment condition as fixed effect, and experiment was a random intercept. Multiple comparison significance values were adjusted by Benjamini–Hochberg.

## Discussion

The human SI has been overlooked not only in gut microbiome research but also as a source and target of microbial therapeutics. Here, we demonstrated how the human SI serves as a target and source of potential microbial therapeutics through isolation and functional screens for enteric hormone production, immunological modulation, and antiviral properties.

The SI microbiome has been difficult to characterize due to its inaccessibility. Many SI samples are collected during endoscopy procedures that are limited to sampling the upper and lower portions of the GI and are typically from patients experiencing GI symptoms. In this study, we collected samples from the SI of human organ donors. There are several drawbacks to microbial sampling from organ donations, such as donors receiving parenteral feeding, intravenous antibiotics, and steroid treatment that may contribute to alterations in the microbial composition of the GI tract^17^. Nevertheless, we identified similar microbial taxa, including *Prevotella*, *Bacteroides*, *Escherichia*, and *Lactobacillus,* as those recovered by ingested devices that sampled distinct sections of the GI in live, healthy adults^56,57^. These parallels reinforce the validity of our findings and underscore the utility of a pill-based sampling approach for high-resolution, site-specific analysis of the microbial and metabolomic profiles of the human SI. Interestingly, this study also report failed library preparation in the device samples, similar to failed preparation in our work, further supporting an extremely low microbial SI biomass in individuals^56^.

Here, we observed strain-specific promotion of secretin and oxytocin in two models **(Fig. 2)**. Lr 103 significantly promoted oxytocin in the adult tissue and not in infant human intestinal organoids while Lr 114 treatment induced secretin in both adult and infant models but only promoted oxytocin in the infant organoid model.**(Fig. 2)**. The selective nature of Lr 103 in the adult jejunal tissue over infant organoid suggests a possible age dependence on the secretion of hormones from this microbe, as there are differences in receptors or cell types in the adult GI epithelium compared to the infant^58,59^.

Nevertheless, the mechanism for stimulation for both hormones remains unclear. Given that secretin is released by low pH, and our supernatants were pH-neutral, the specific stimulation of secretin by Lr 103 and Lf 114 indicates an unknown microbially-produced molecule stimulating secretin. The secretion of secretin and oxytocin from Lr 103 and Lf 114 aligns with the findings in Danhof et al., where *L. reuteri* 6475 promoted the release of secretin, which stimulated the release of oxytocin the GI tract^30^. However, we additionally observed a secretin-independent secretion of oxytocin from *L. reuteri* 6475 in the infant models. Unlike Danhof et al., our study cultured *L. reuteri* 6475 in DMEM for organoid experimentation **(Fig. 2C,2D)**. The substantially lower secretin levels detected (22 pg/mL, **Fig. 2C** compared to 75 pg/mL^30^), suggest either altered *L. reuteri* 6475 metabolism in DMEM or a secretin-independent mechanism of oxytocin release^30^. Microbial short-chain fatty acid production^61^, secreted proteins^62^, and modulation of host hormonal gene expression^76^ have all been previously found to impact enteric hormone secretion and provide initial avenues to explore secretin-independent promotion of oxytocin release in the human gut. Our study highlights the potential for SI microbes to modulate critical metabolic-related hormones that could have implications in satiety, wound healing, and gut-brain interactions^62, 63,64^.

*Lactobacillaceae* are also known for their modulation of the immune system^42^ ^65,40^ ^66^ ^67^ ^49^ ^68^ ^69^. While further research is required to identify how the candidate microbial therapeutics function *in vivo* with the host immune response and complex microbiome, we can speculate how the potential microbial therapeutics could be applied. The pro-inflammatory effects of Ls 112 could function as an adjuvant in conjunction with immunotherapy or be applied as a therapy directly within a tumor to stimulate a specific immune response **(Fig. 3)**^70^. In disease states that would require an anti-inflammatory therapy, such as in IBD, enteric infection, or other GI inflammation, La 180 and Lf 114 could be used to dampen and modulate host response. Specifically, in TNF-driven disease such as IBD, La 180 and Lf 114 could function as specific drivers of anti-inflammation. Future mechanistic-focused and *in vivo* studies could glean more information on the strength of candidacy of these SI isolates.

IFN-λ secretion from infant organoids was revealed to be via TLR3 after and dependent on a polyI:C prime with bacterial supernatant treatment **(Fig 4**, **Fig 5)**. Previous research has described *Lactobacillaceae* promotion of the secretion or the expression of IFN-λ in multiple models with one paper specifically reporting dsRNA from *Lactobacillaceae* activating TLR3 and promoting secretion of IFN-β^71, 72^ ^51, 52^. Interestingly, bacterial supernatant alone did not promote IFN-λ secretion in our studies, despite our results supporting that Lr 103 secretes RNA which induces IFN-λ through TLR3 **(Fig.4**, **Fig. 5)**. We speculate that since TLR3 is typically expressed at low levels at homeostasis, the highly concentrated polyI:C prime likely provides a more potent stimulus than bacterial supernatant alone^15,73^. The polyI:C prime indirectly promotes TLR3 by binding TLR3, which promotes downstream expression of interferon-stimulated genes, including TLR3, enabling multimeric TLR3 dimerization along longer dsRNA that elicits a stronger IFN-λ response^74,75,76^. Lastly, organoids are less responsive to bacterial stimulation when cultured in antioxidant-rich differentiation media^73^. While we did not identify the mechanism behind how Lr 103 is delivering the RNA to the organoids, one way may be through extracellular vesicles, which have been described as having diverse RNAs packaged inside that can impact host function^77^. In addition to the antiviral activity, IFN-λ has also been associated with reduced inflammation in mouse models of arthritis, possesses anti-tumor activity, and inhibits proliferation^53^. Lr 103 could be further developed as a potential MT candidate to reduce inflammation in arthritis, other autoimmune diseases, and as an adjuvant to chemotherapy to reduce proliferation of cancerous cells.

IFN-λ is important in the early stages of rotavirus infection in a mouse model of rotavirus infection^47^. A mild effect of IFN-λ in rotavirus replication inhibition is in line with our findings, where Lr 103 and La 180 mildly inhibited RV1 replication in infant organoids and were independent of IFN-λ. We posit that this enhanced TLR3 stimulation from the presence of Lr 103 during infection prompts a fast, antiviral activity in the organoids, with an IFN-λ effect that is damped by interferon antagonism by rotavirus.

IFN-λ has been demonstrated to inhibit wild-type rotavirus replication in mice and adult organoids ^55,47^. Following the observation that Lr 103 induced IFN-λ secretion following a polyI:C prime **(Fig. 4**, **Fig. 5)**, and because the infant organoids used in our study were established from infants age-eligible for oral rotavirus vaccines, we evaluated the effect of Lr 103 on oral rotavirus vaccine replication. A significant reduction of oral rotavirus vaccine replication was observed in infant organoids. These results suggest that factors like the age of recipients and vaccination schedules should be considered when using SI microbes as microbial therapeutics. Interestingly, other studies have evaluated lactobacillus strains to improve immune response to live attenuated ^78^and inactivated^79, 80^ influenza vaccines and oral rotavirus vaccines^81^. The inhibitory effect of Lr 103 on RV1 replication observed in our study also indicates that different lactobacillus strains may have differing effects on vaccine take and could be explored in infectious rotavirus strains in the future.

In summation, our study has provided evidence of a potential pipeline for isolation of SI microbes through pre-clinical evidence of potential translational impact on human health. Future directions include mechanistic investigations of these beneficial effects, *in vivo* characterization of therapeutic potential, and progression of promising strains through the FDA approval pipeline for microbial drug development.

## Acknowledgements

We would like to thank the organ donors and their families for their generous donation and support for our science. We also acknowledge Drs. Sara Di Rienzi, Heather Danhof, Micah Forshee, Katherine Wozniak, and Alexa Corker for their feedback during project design and manuscript preparation. The authors would also like to thank the sequencing and project management contributions completed by Mathew Ross and Juwan Cormier at the CMMR.

## Author Contributions

E.J.N. contributed to study design, data acquisition and analysis, manuscript preparation, and manuscript editing and statistical analysis. L.N.S performed rotavirus infection assays, provided feedback on study design, and edited the manuscript. A.K.B.A completed identification of microbes by whole genome analysis. S.R. provided rotavirus vaccine strains, feedback on the study design, critical reviews of the manuscript, and approval of the article. R.A.B. contributed to concept, study design, manuscript editing, critical review, approval of the article, and funding acquisition.

---

**Supplementary Figure 1.**
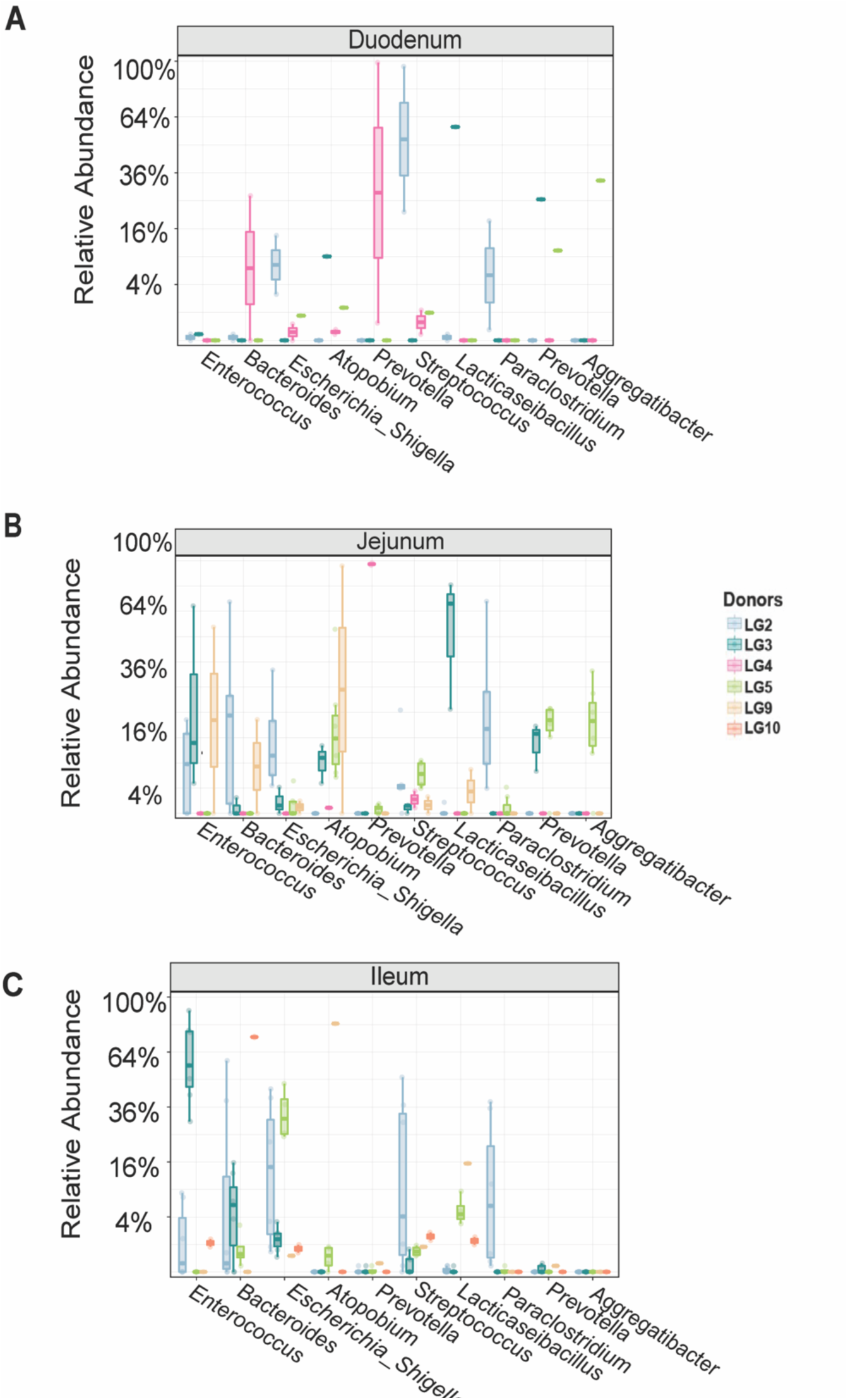
**Cultivation of SI microbial community from organ donors**. The top 10 taxa are presented in the duodenum (A), jejunum (B), and ileum (C) from cultivation-independent (sequence-only) samples from donors 2-5 and 8 using V4 16S rRNA sequencing.

**Supplementary Figure 2.**
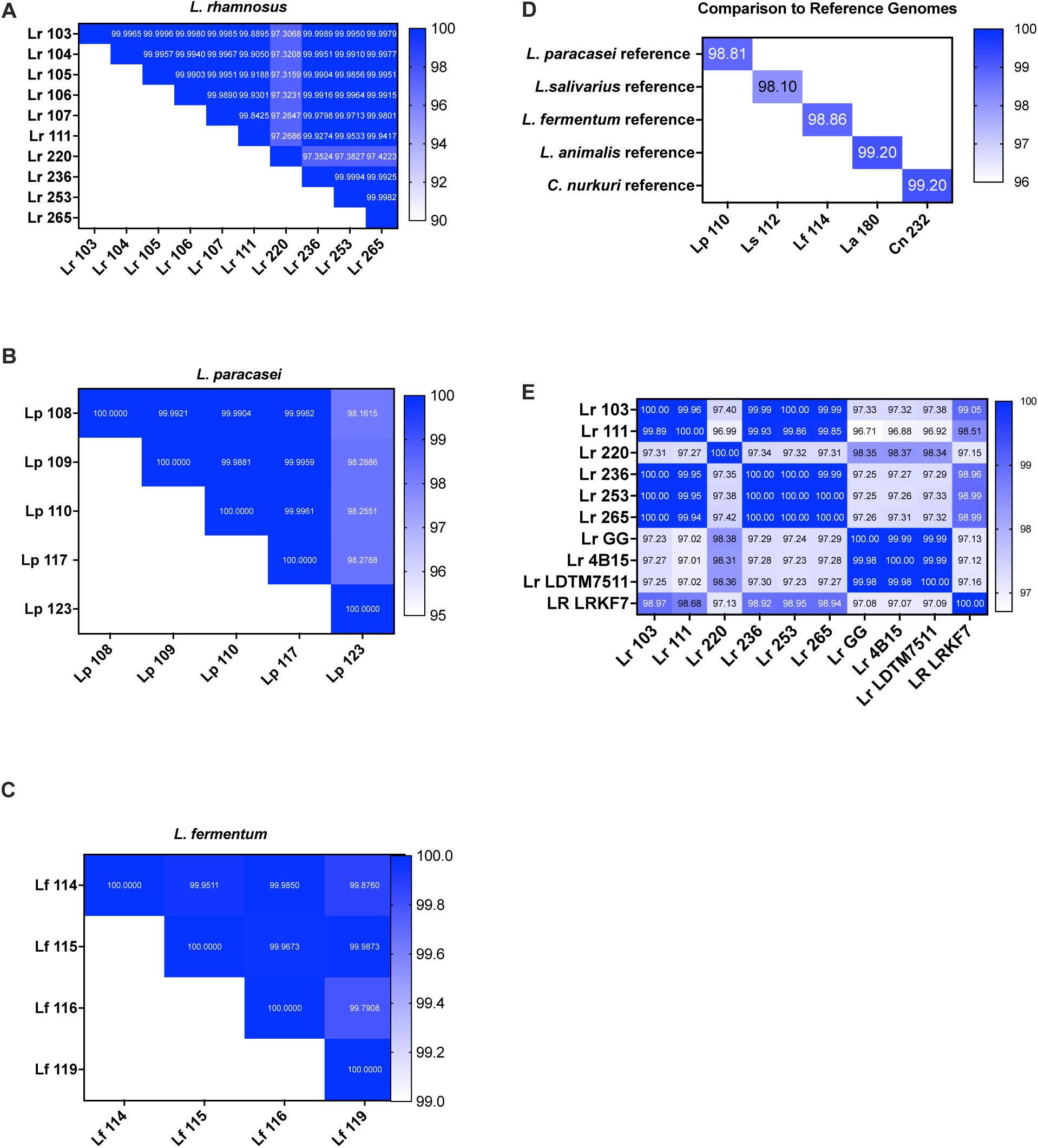
Average nucleotide identity analysis of SI *Lactobacillaceae*. The average nucleotide identity (ANI) pairwise analysis was completed on samples with multiple isolates that were identified as *L. rhamnosus* **(A)***, L. paracasei* **(B)**, and *L. fermentum* **(C)** by whole-genome sequencing. **(D)** SI isolates were compared by ANI to their reference genomes from NCBI. **(E)** *L. rhamnosus* strains isolated in our study were compared by ANI to commercial *L. rhamnosus* products.

**Supplementary Figure 3.**
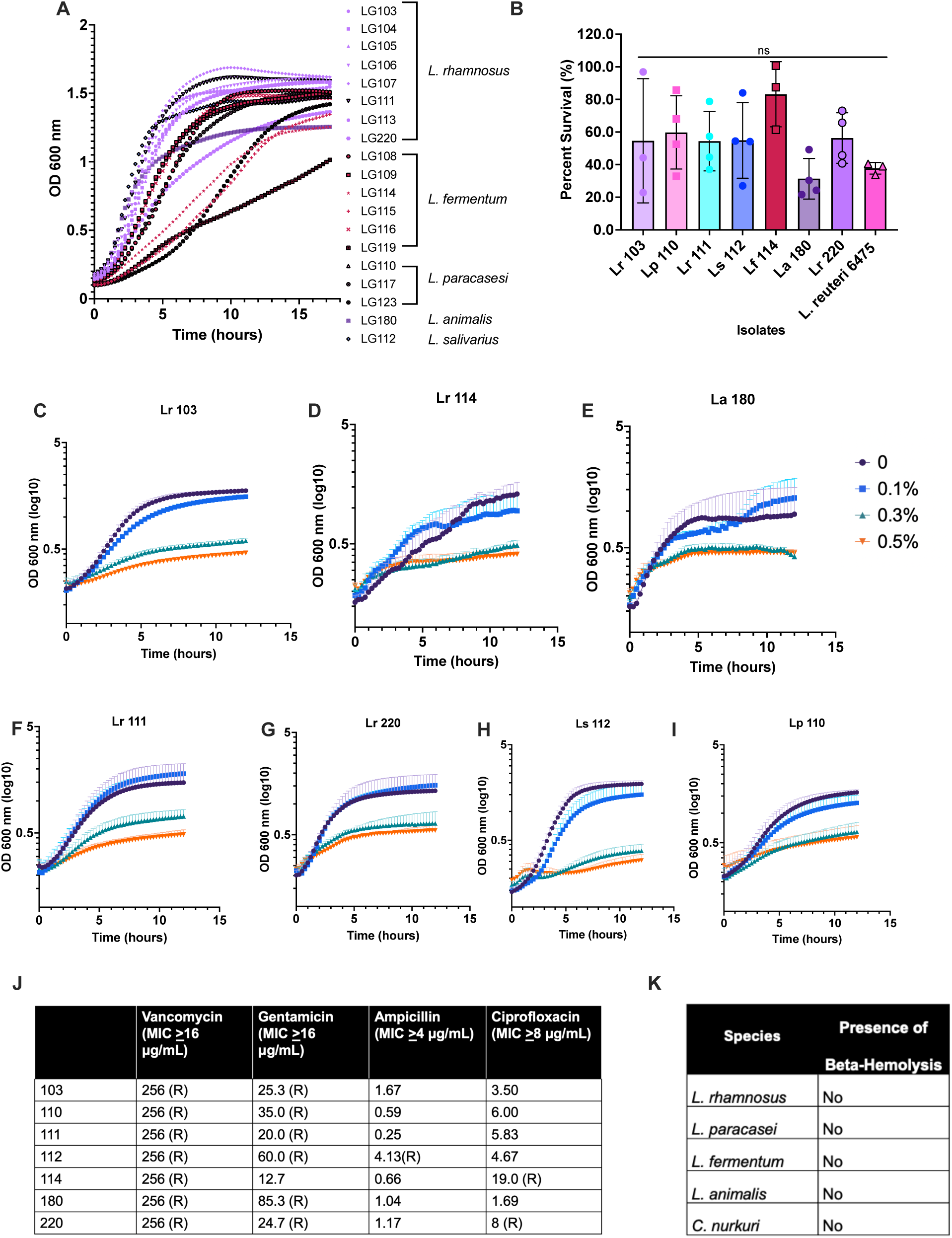
Isolates withstand GI-like stress and exhibit safety indicators. **(A)** Growth curves in MRS were determined by measuring OD_600_ nm every 20 minutes in a 96-well plate with calculated doubling times (n=3). **(B)** Strains were exposed to pH 2 MRS for 2 hours to test for acid resistance. Isolates were plated at time 0 and time 2 hours in treatment; CFU/ mL was calculated from those two time points to determine survivability n=3 (time 2/ time 0). One-way ANOVA was completed with multiple comparison corrections, and strains were not statistically significant when compared to *L. reuteri* 6475. **(C-I)** Bile resistance in 0, 0.1%, 0.3%, and 0.5% was measured by OD 600 nm every 20 minutes in 96-well plates over 12 hours to record growth for each isolate (n=3). **(L)** Antibiotic resistance was measured using MIC test strips, and R indicates resistance to the antibiotic. **(J)** Strains were both spotted from overnight cultures or streaked from a colony onto Columbia blood agar plates. **(K)** Beta-hemolysis was determined by complete clearance in the agar (n=2).

**Supplementary Figure 4.**
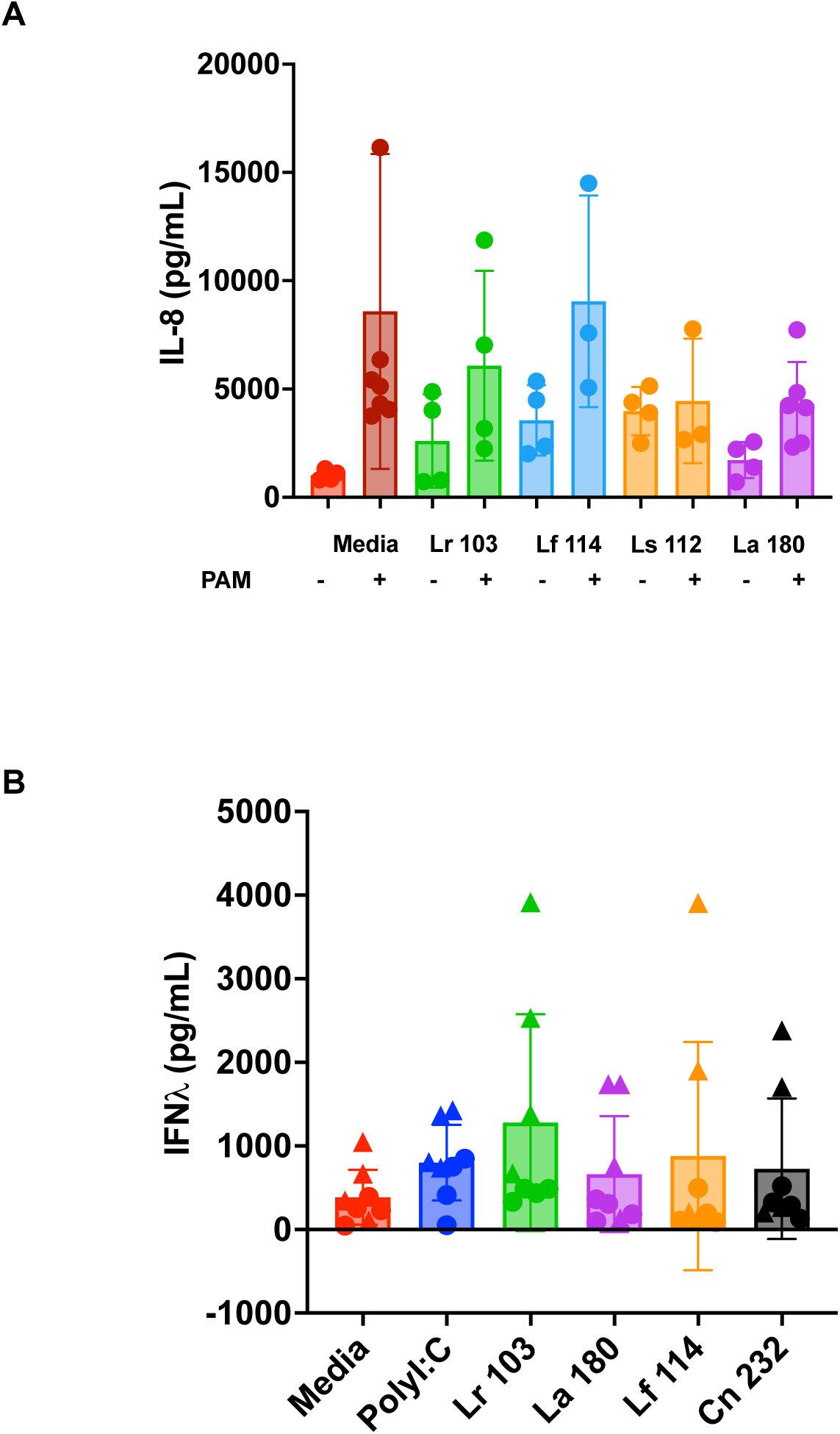
**Cytokine response from treatment of cell-free Si bacterial supernatant**. **(A)** Bacterial supernatants were added to THP-1 cells or primed with TLR-2 agonist, and IL-8 was by ELISA (n=4-6). Statistics were calculated by a linear model with treatment as a fixed effect with each measurement treated as an independent observation. Multiple comparison significance values were adjusted by Benjamini–Hochberg. Significances are designated by * indicating p ≤0.05, ** indicating p ≤0.01, and *** indicating p ≤0.001. **(B)** Small intestinal organoid monolayers were primed with 50 ug/mL of polyI:C, then treated with 4 different bacterial supernatants to screen for IFN-λ activity (n=4). Shapes represent J1006 (circle) and IL1002 (triangle).

**Supplementary Figure 5.**
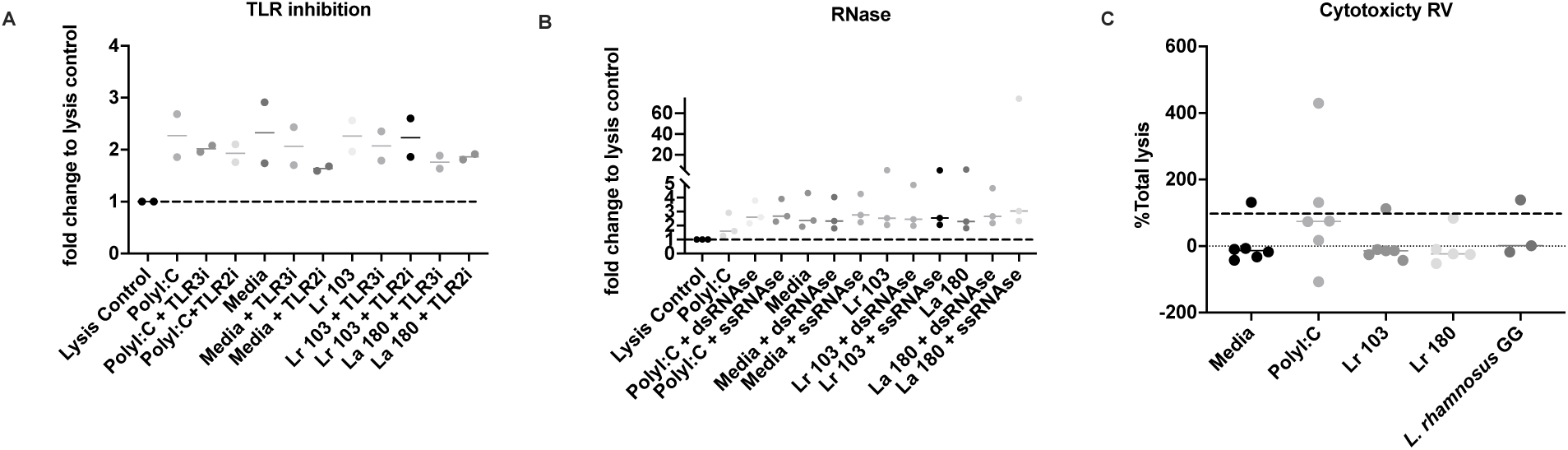
Cell viability after experimentation. Presto-blue assays were completed to determine the viability of the organoids after the live-addition test **(A)**, addition of dsRNAse or ssRNAse **(B)**, and TLR inhibitors **(C)**. Cytotoxicity assays were done after incubation with RV1 **(D)**.

**Table S1.**
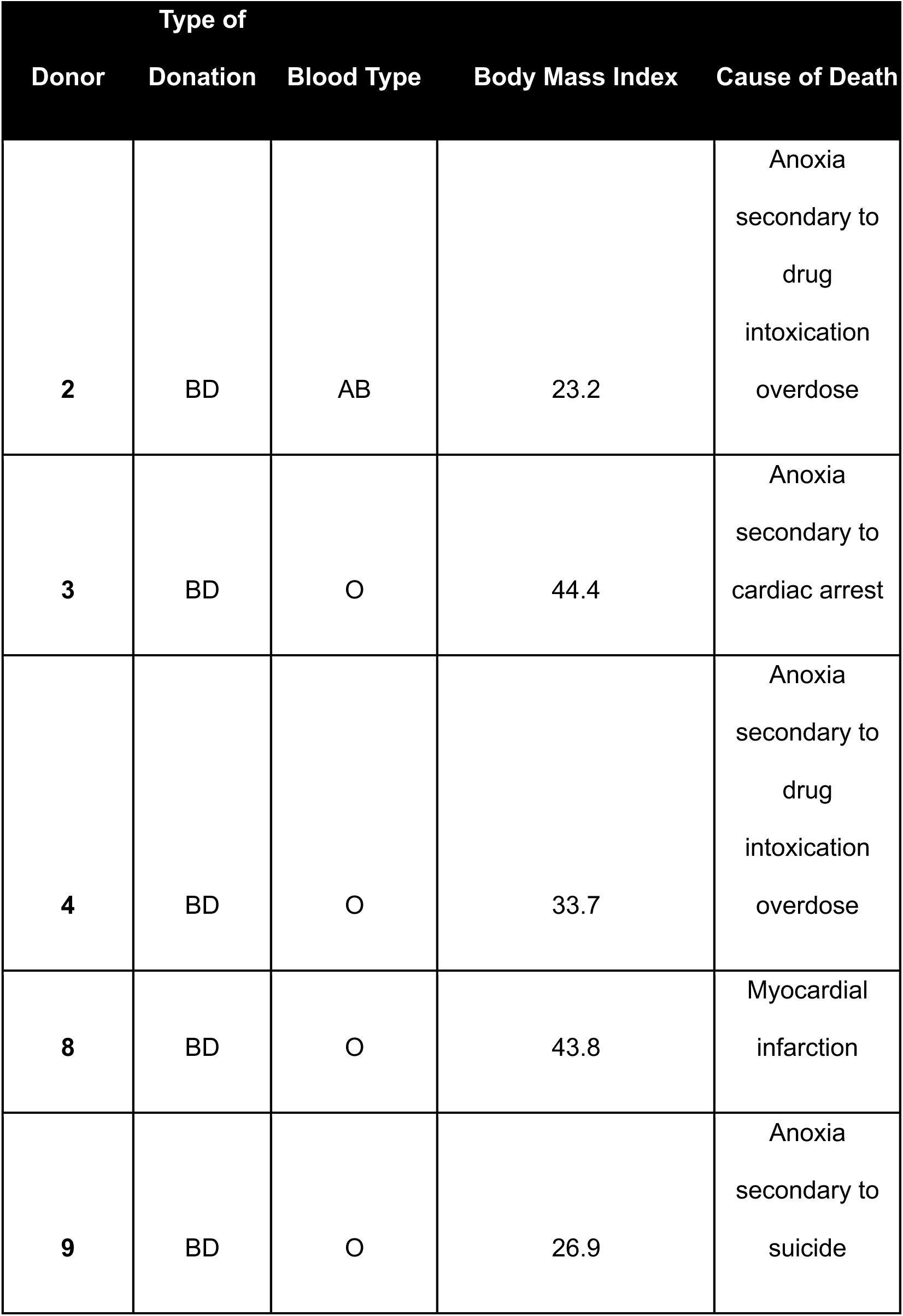
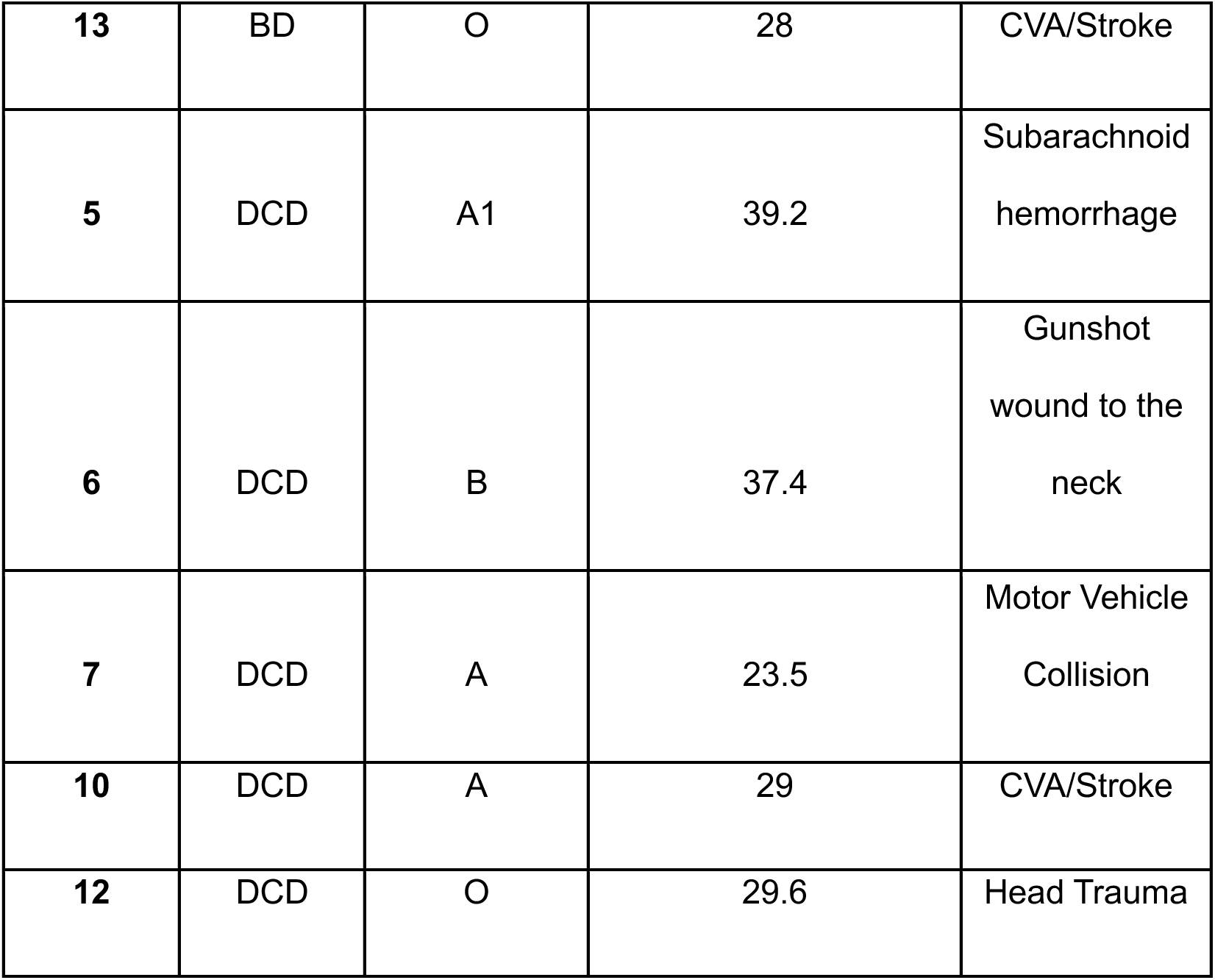
Demographic info from organ donors. Extended information on the donors, such as blood type, body mass index, and cause of death. Adapted from Nachman et al (2025).

**Table S2.**
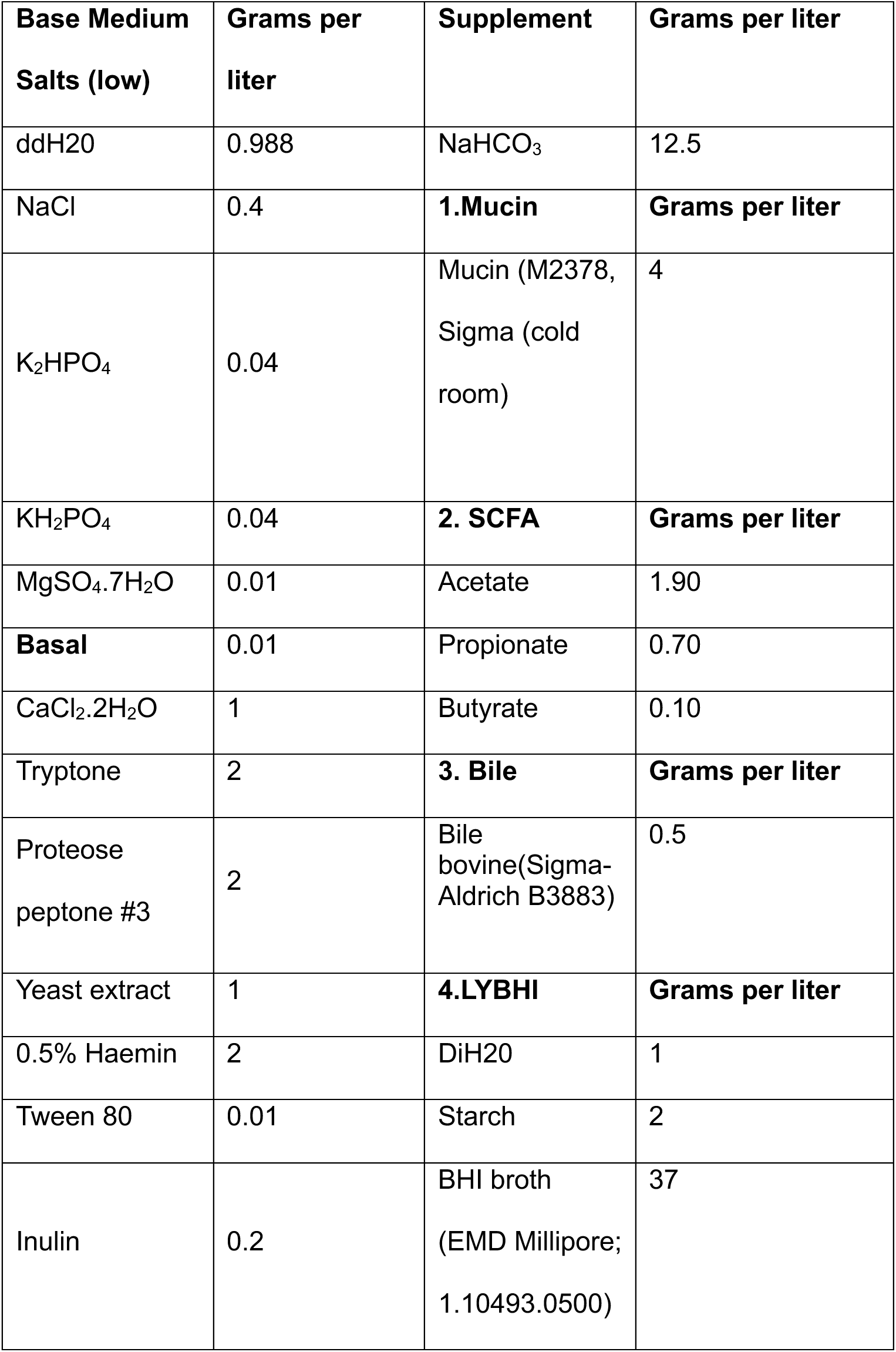
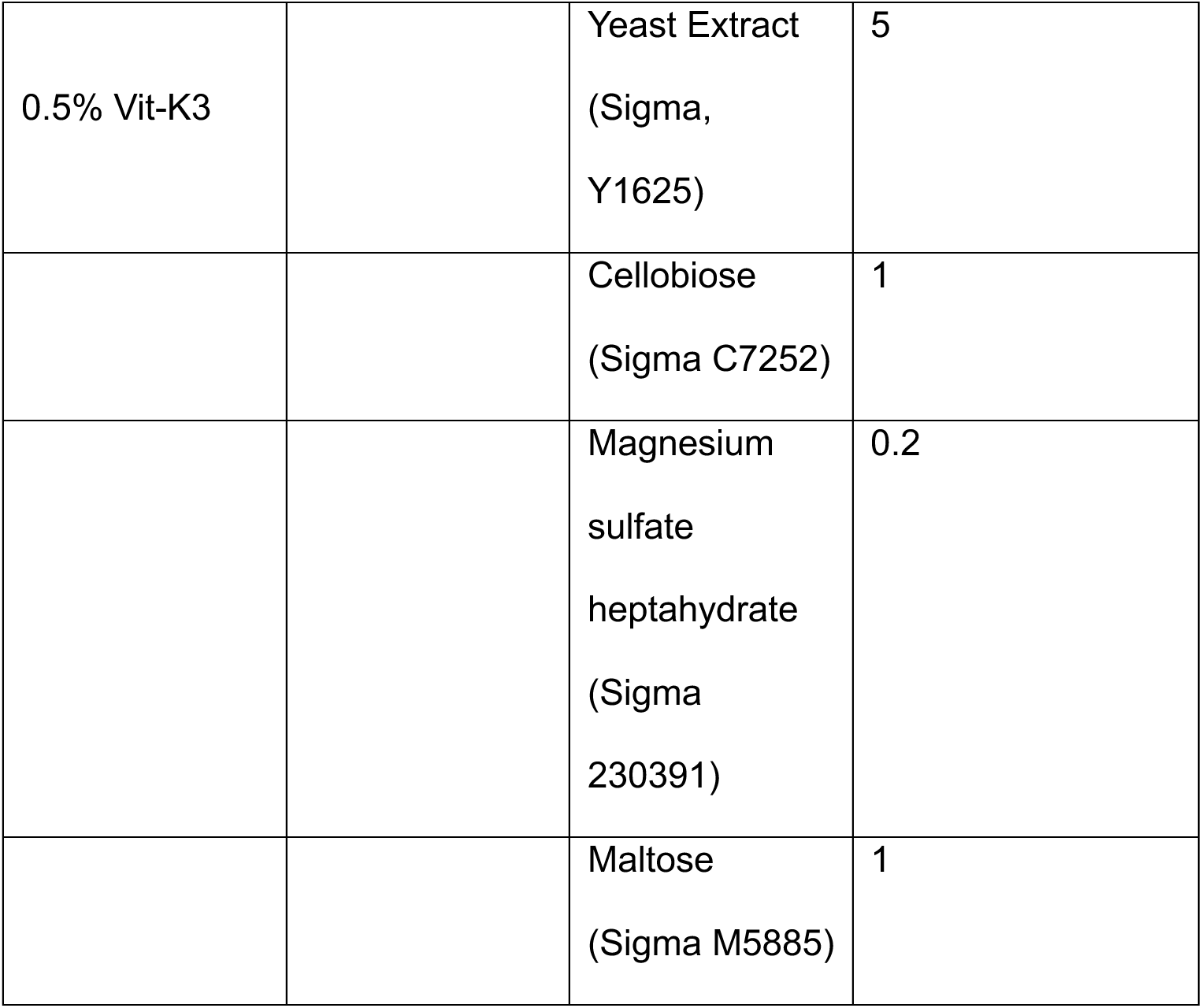
Media composition for GIT microbial isolation.

**Table S3.** Infant organoid demographic information.

| **Name** | **Sex** | **Age at Tissue Retrieval (months)** | **Intestinal Section** |
| --- | --- | --- | --- |
| J1005 | Female | 2 | Jejunum |
| J1006 | Female | 3 | Jejunum |
| J1009 | Male | 5 | Jejunum |
| II1002 | Male | 24 | Ileum |
| IL1004 | Male | 3 | Ileum |
| IL1010 | Male | 5 | Ileum |
| IL1013 | Female | 3 | Ileum |

